# Peptide Pattern Recognition for high-throughput protein sequence analysis and clustering

**DOI:** 10.1101/181917

**Authors:** Peter Kamp Busk

## Abstract

Large collections of protein sequences with divergent sequences are tedious to analyze for understanding their phylogenetic or structure-function relation. Peptide Pattern Recognition is an algorithm that was developed to facilitate this task but the previous version does only allow a limited number of sequences as input.

I implemented Peptide Pattern Recognition as a multithread software designed to handle large numbers of sequences and perform analysis in a reasonable time frame. Benchmarking showed that the new implementation of Peptide Pattern Recognition is twenty times faster than the previous implementation on a small protein collection with 673 MAP kinase sequences. In addition, the new implementation could analyze a large protein collection with 48,570 Glycosyl Transferase family 20 sequences without reaching its upper limit on a desktop computer.

Peptide Pattern Recognition is a useful software for providing comprehensive groups of related sequences from large protein sequence collections.

## Introduction

Peptide Pattern Recognition (PPR) is a non-alignment-based method for analyzing large number of divergent protein sequences (Busk and Lange, 2013). The method consists of identifying a defined number of short sequences that are conserved in a group of protein sequences each containing more than a threshold number of the short, conserved sequences in their amino acid sequence. Hence, the output of PPR consists of groups of proteins with corresponding groups of short sequences that are conserved in the protein sequences.

The proteins in each PPR group have the same tertiary fold (Busk *et al*., 2017) and often share functional features such as similar or same function; e.g. PPR predicts the function of carbohydrate-active enzymes with higher precision than any other method (Busk and Lange, 2013). Due to these features PPR has been used to provide an overview of the sequence variation of new enzyme families to pinpoint conserved motifs in the enzymes and find sequence features related to function (Busk and Lange, 2015; Agger *et al*., 2017). Moreover, the short, conserved peptides can be used as a fast tool for finding enzymes similar to a PPR group in genomes, transcriptomes and other sequence data (Bech *et al*., 2014; Busk *et al*., 2014; Huang *et al*., 2014; Busk *et al*., 2017; Wilkens *et al*., 2017).

Recently, PPR was used to divide the integral membrane fatty acid desaturase family into groups of sequences that could be used for phylogenetic analysis (Wilding *et al*., 2017). However, only a limited number of the sequences could be grouped due to low processing capacity of the previously published PPR software.

Here, I present an updated PPR package (PPR version 2) capable of grouping at least 48,570 sequences in 50 hours on a desktop computer.

## Implementation

The PPR algorithm (Busk and Lange, 2013) was implemented in an updated PPR package to boost the number of sequences that can be processed and to increase the speed of analysis. PPR version 2 is provided as source code with the possibility to modify all relevant parameters including selecting a range of parameters to screen, set up automatic screening of several protein families or both.

The input and output files are defined directly in the source code or by providing them as a variable when running PPR. The same goes for the most important parameters for PPR analysis: the length of the conserved sequences (peptide length), number of conserved sequences (limit) and number of conserved sequences in each protein (cut off) (Busk and Lange, 2013). Finally, a number of parameters that are usually not changed from run to run are defined in the source code.

The result of a PPR analysis consist of a number of files containing groups of protein sequences and a number of files containing lists of short sequences (peptides) that are highly conserved in the protein groups.

The file {input name}_classification_overview.txt contains an overview of the number of proteins in each group and the file {input name}_conserved_peptides.txt contains a short list of the conserved peptides and their frequency for each group. This list is suitable for annotation of new proteins to the groups, e.g.; as an input to the applications classify proteins (Busk and Lange, 2013), https://sourceforge.net/projects/shocopop/files/ or Homology to Peptide Patterns (Hotpep) (Busk *et al*., 2014, 2017).

## Results

Analysis of different sequence collections (see Supplemental Files) showed that PPR version 2 was faster than the previously published PPR software (old PPR) (Busk and Lange, 2013) for the small sequence collections with 104 Glycosyl Hydrolase family 117 (GH117) enzymes and for 637 MAP kinases (MAPK)(Table 1). The old PPR crashed when handling 1253 Zinc-finger proteins (ZNF).

**Table 1:** Benchmarking and performance of PPR version 2

| Proteins | Sequences | Analysis time<br>old PPR | Analysis time<br>PPR version 2 | Groups | Percent identity |
| --- | --- | --- | --- | --- | --- |
| GH117 | 104 | 34 s | 4 s | 5 | 18 - 36 |
| MAPK | 673 | 1680 s | 72 s | 16 | 8 – 43 |
| ZNF | 1,253 | na | 170 s | 8 | 10 – 56 |
| P450 | 1,355 | na | 316 s | 48 | 8 – 44 |
| GH20 | 2,343 | na | 1432 s | 57 | 8 – 43 |
| GT2 | 48,570 | na | 50 h | 234 | 0 – 52 |

Analysis time was measured on a desktop computer (IntelR Core i7-2600 CPU @ 3.40 GHz, 16 GB RAM). Percent identity between highest scoring sequence in each group was calculated with MUSCLE (Edgar, 2004). “na”: Not analyzable.

In contrast PPR version 2 could handle this and larger sequence collections up to 48,570 Glycosyl Transferase family 2 (GT2) sequences (Table 1). No further testing of larger families was performed as most families are smaller than 50,000 members. Exceptionally large families can be handled by using a more powerful hardware setup.

The diversity of sequences within each PPR group is large as measured by MUSCLE pairwise alignment (Edgar, 2004) of the highest scoring protein in each group (Table 1).

## Conclusion

PPR version 2 is a fast method for dividing large families of protein sequences into related groups to facilitate phylogenetic analysis (Wilding *et al*., 2017) and functional sequence analysis (Agger *et al*., 2017; Busk and Lange, 2015, 2013). The lists of conserved, short sequences provided for each group can be used for structure-function investigations and for annotation of proteins e.g. predicted protein coding sequences from a genome with the Hotpep program (Busk *et al*., 2014).

The software can be used on a desktop computer and is available as source code for modification and improvement.

## Availability

Peptide Pattern Recognition is available at https://sourceforge.net/projects/peptide-pattern-recognition/

## Acknowledgements

None.

